## Additional File 1_Supplementary methods for "Plant Growth Promoting fungal endophyte *Colletotrichum tofieldiae* Ct0861 reduces mycotoxigenic *Aspergillus* fungi in maize grains"

### 1. Soil physical-chemical properties in the control and Ct0861 plots.[1]

| PROPERTIES | | | UNIT | BEFORE EXPERIMENT[2] | | AFTER EXPERIMENT[3] | | Standard[4] | Method/Assay |
|---|---|---|---|---|---|---|---|---|---|
| | | | | Control | Ct0861 | Control | Ct0861 | | |
| SOIL TEXTURE | Sand (2-0.05 mm) | | %(w/w) | 54 | 50 | 70 | 68 | - | Bouyoucos densimeter method |
| | Silt | | %(w/w) | 12 | 16 | 12 | 10 | - | Bouyoucos densimeter method |
| | Clay | | %(w/w) | 34 | 34 | 18 | 22 | - | Bouyoucos densimeter method |
| | Bulk density | | g/cc | 1.391 | 1.396 | 1.468 | 1.46 | - | Calculation |
| SALINITY | Electrical conductivity (25ºC) liquid ext. 1/5 (w/v) | | mS/cm | 0.303 | 0.379 | 0.693 | 0.635 | UNE-EN 13038 | Electrical conductivity by conductimetry |
| | Sol. chloride in liquid ext. 1/5 (v/v) | Cl | meq/100g | 0.159 | 0.194 | 0.72 | 0.53 | UNE 77308 | Electrical conductivity by conductimetry |
| | Sol. sulfate in liquid ext. 1/5 (w/v) | Gypsum | %(w/w) | 0.0294 | 0.0493 | 0.064 | 0.0701 | UNE 77308 | Electrical conductivity by conductimetry |
| | Available Sodium | Na | meq/100g | 0.62 | 0.64 | 1.12 | 1.13 | UNE-EN ISO 11260 | Extractable metals with barium chloride and triethanolamine (TEA) at pH 8.2 by inductively coupled plasma emission spectroscopy (ICP/AES) |
| SOIL REACTIVITY | pH in KCl 1M extract 1/2 (v/v) | | pH Units | 7.79 | 7.81 | 7.77 | 7.74 | UNE-EN 13037 | pH by potenciometry |
| | Total limestone | $CaCO_3$ | %(w/w) | 33.21 | 28.37 | 29.21 | 4.23 | | Bernard calcimeter method |
| | Active limestone | $CaCO_3$ | %(w/w) | 10.3 | 10.17 | 9.34 | 2.829 | | Ammonium oxalate extraction |

[1]Approximately 500 g soil material from the top 30 cm layer were collected and pooled from the four plots of each treatment (control or Ct0861).
[2] Samples were taken before the initiation of the experiment and the fertilization of the soil.
[3] Samples were taken after the harvest of the experiment on soil that had been fertilized and cultivated with maize.
[4] UNE: Spanish Standarization Organism; ISO: International Organization for Standarization; SSIR: Survey Soil Laboratory Information Manual (United States Department of Agriculture)

| PROPERTIES | | | UNIT | BEFORE EXPERIMENT[2] | | AFTER EXPERIMENT[3] | | Standard [4] | Method/Assay |
|---|---|---|---|---|---|---|---|---|---|
| | | | | Control | Ct0861 | Control | Ct0861 | | |
| ORGANIC MATTER | Total organic matter | | %(w/w) | 2.28 | 2.12 | 2.03 | 1.96 | SSIR 42, Method (6A1) | Organic matter by volumetry (potentiometric titration) |
| | Total Organic Carbon | C | %(w/w) | 1.323 | 1.231 | 1.175 | 1.138 | SSIR 42, Method (6A1) | Organic matter by volumetry (potentiometric titration) |
| | Carbon/Nitrogen rate | C/N | | 8.43 | 7.89 | 7.08 | 6.98 | | Calculation, Organic C/Total N |
| PRIMARY MACRONUTRIENTS | Total Nitrogen | N | %(w/w) | 0.157 | 0.156 | 0.166 | 0.163 | UNE-EN 16168 | Total nitrogen by thermal conductivity (Dumas method) |
| | Nitric nitrogen sol. In liquid extract 1/5 (w/v) | N | mg/kg | 34.8 | 34.8 | 63.1 | 61.3 | UNE-EN 10304-1 | Soluble anions in aqueous extract by ion chromatography with electrical conductivity detector |
| | Available Phosphorus | P | mg/kg | 138 | 134 | 119 | 113 | ISO 22036 | Phosphorus by inductively coupled plasma emission spectroscopy (ICP/AES) (Olsen method) |
| | Available Potassium | K | meq/100 g | 2.33 | 2.11 | 1.81 | 1.97 | UNE-EN ISO 11260 | Extractable metals with barium chloride and triethanolamine (TEA) at pH 8.2 by inductively coupled plasma emission spectroscopy (ICP/AES) |
| SECONDARY MACRONUTRIENTS | Available Calcium | Ca | meq/100g | 10.5 | 9.9 | 10 | 9.7 | UNE-EN ISO 11260 | Extractable metals with barium chloride and triethanolamine (TEA) at pH 8.2 by inductively coupled plasma emission spectroscopy (ICP/AES) |

| PROPERTIES | | | UNIT | BEFORE EXPERIMENT[2] | | AFTER EXPERIMENT[3] | | Standard [4] | Method/Assay |
|---|---|---|---|---|---|---|---|---|---|
| | | | | Control | Ct0861 | Control | Ct0861 | | |
| | Available Magnesium | Mg | meq/100g | 2.8 | 2.73 | 2.85 | 2.83 | UNE-EN ISO 11260 | Extractable metals with barium chloride and triethanolamine (TEA) at pH 8.2 by inductively coupled plasma emission spectroscopy (ICP/AES) |
| MICRONUTRIENTS | Available Iron | Fe | mg/Kg | 9.3 | 7.5 | 6.72 | 6.32 | UNE-EN 13651 | Extractable metals with diethylenetriamine pentaacetic acid (DTPA), calcium chloride and TEA at pH 7,3 by atomic emission spectroscopy with inductive coupled plasma (ICP/AES) |
| | Available Manganese | Mn | mg/Kg | 15.7 | 13.5 | 9.5 | 8.9 | UNE-EN 13651 | Extractable metals with diethylenetriamine pentaacetic acid (DTPA), calcium chloride and TEA at pH 7,3 by atomic emission spectroscopy with inductive coupled plasma (ICP/AES) |
| | Available Zinc | Zn | mg/Kg | 11.5 | 10.2 | 10.1 | 9.6 | UNE-EN 13651 | Extractable metals with diethylenetriamine pentaacetic acid (DTPA), calcium chloride and TEA at pH 7,3 by atomic emission spectroscopy with inductive coupled plasma (ICP/AES) |
| | Available Copper | Cu | mg/Kg | 3.95 | 3.15 | 3.82 | 3.28 | UNE-EN 13651 | Extractable metals with diethylenetriamine pentaacetic |

| PROPERTIES | | | UNIT | BEFORE EXPERIMENT[2] | | AFTER EXPERIMENT[3] | | Standard [4] | Method/Assay |
|---|---|---|---|---|---|---|---|---|---|
| | | | | Control | Ct0861 | Control | Ct0861 | | |
| | | | | | | | | | acid (DTPA), calcium chloride and TEA at pH 7,3 by atomic emission spectroscopy with inductive coupled plasma (ICP/AES) |
| | Available Boron | B | mg/Kg | 1.13 | 1.18 | 0.97 | 1.03 | | Extractable metals in aquaeus extraction by atomic emission spectroscopy with inductive coupled plasma (ICP/AES) |
| AVAILABLE CATIONS: RELATIVE PROPORTIONS | Exchangeable Sodium Percentage (ESP) | | % | 3.8 | 4.2 | 7.1 | 7.2 | | Calculation |
| | Potassium Relative Proportion | | % | 14.3 | 13.7 | 11.5 | 12.6 | | Calculation |
| | Calcium Relative Proportion | | % | 64.7 | 64.5 | 63.4 | 62 | | Calculation |
| | Magnesium Relative Propotion | | % | 17.2 | 17.7 | 18.1 | 18.2 | | Calculation |
| AVAILABLE CATIONS: INTERACTIONS | Ratio Calcium/Magnesium | Ca/Mg | % | 3.77 | 3.64 | 3.51 | 3.41 | | Calculation |
| | Ratio Potassium/Magnesiu m | K/Mg | % | 0.83 | 0.77 | 0.64 | 0.7 | | Calculation |

### 2. Irrigation program for open-field maize plants.

| | CROP ESTABLISHMENT | VEGETATIVE DEVELOPMENT | | REPRODUCTIVE STAGE | SENESCENCE | | |
|---|---|---|---|---|---|---|---|
| | | Stage 1 | Stage 2 | | Stage 1 | Stage 2 | Stage 3 |
| No. days per period | 20 | 17 | 18 | 40 | 10 | 10 | 10 |
| Watering regime until | 28/05/18 | 14/06/18 | 02/07/18 | 11/08/18 | 21/08/18 | 31/08/18 | 10/09/18 |
| Kc[1] | 0.4 | 0.79 | 1.20 | 1.20 | 0.92 | 0.63 | 0.35 |
| Eto (mm/day) | 4.80 | 5.25 | 5.72 | 5.55 | 5.16 | 4.73 | 4.32 |
| Rainfall (mm/day) | 0.30 | 0.14 | 0.20 | 0.10 | 0.29 | 0.35 | 0.62 |
| Effective rainfall (mm/day) | 0.1 | 0.0 | 0.0 | 0.0 | 0.1 | 0.1 | 0.3 |
| ETC (mm/day) | 1.92 | 4.14 | 6.86 | 6.66 | 4.73 | 2.99 | 1.51 |
| GIWR (mm/day) | 1.82 | 4.14 | 6.84 | 6.66 | 4.64 | 2.86 | 1.17 |
| NIWR (mm/day) | 1.80 | 4.10 | 6.77 | 6.60 | 4.59 | 2.83 | 1.16 |
| TIWR (mm/day) | 2.37 | 5.39 | 8.91 | 8.68 | 6.04 | 3.72 | 1.53 |
| Watering time (h/day) | 0.51 | 1.16 | 1.91 | 1.86 | 1.29 | 0.80 | 0.33 |
| Watering time in OW (min/day) | 30 | 69 | 115 | 112 | 78 | 48 | 20 |

| | |
|---|---|
| Eto | Reference evapotranspiration |
| ETC | Reference crop evapotranspiration |
| GIWR | Gross irrigation water requirements |
| NIWR | Net irrigation water requirements |
| TIWR | Total irrigation water requirements |

Calculations were made on long-term data from the closest climate station, located in San Javier (37° 47' 20'' N, 0° 48' 12'' O; IMIDA 2018).

[1]Corn Kc was adapted from CROPWAT (version 8.0) and seed breeder information.

### 3. Processing of soil and plant material until DNA extraction

BULK SOIL COMPARTMENT: Approximately 0.5 L of soil material from the top 30 cm surrounding layer of each plant were collected in a plastic zip bag. Then, 100 mg were directly transferred to a 2 mL Eppendorf tube with lysis matrix from FastDNA™ Spin Kit for Soil (MP Biomedicals).

RHIZOSPHERE COMPARTMENT: Roots were separated from the rest of the plant and shaken to remove the excess of soil. Roots were washed twice in two 500 mL plastic recipients with 100 mL of sterile distilled water to remove attached soil particles. The 200 mL of washing water (containing soil particles) from the same plant, were mixed and 30 mL were transferred to a 50 mL falcon. Falcon tubes were centrifuged at 4,000xg for 15 min. After centrifugation, 95% of supernatant was discarded. Pellet was resuspended into the remaining supernatant. Using a cut P1000 tip, 300 µL of the mixture were transferred to a 2 mL tube with lysis matrix from FastDNA™ Spin Kit for Soil (MP Biomedicals).

ROOT COMPARTMENT: Roots were gently washed with tap water in order to remove any remaining soil residue. Roots were cut into small sections with pruning shears, which were disinfected with a 20% bleach solution between samples. Root sections were wrapped in aluminum foil for their storage.

LEAF COMPARTMENT: Four circles of ~1 cm of diameter were collected from the youngest unfolded leaf of each plant and introduced in a 2 mL Eppendorf tube filled with sterile glass beads (2.7 mm Ø, Carl Roth GmbH & Co.).

GRAIN COMPARTMENT: Grain samples were only collected for 4MPS. Mature cobs from the same plant were manually shelled, grains were pooled and transferred to 15 mL falcon.

All samples were kept at -80ºC until DNA extraction.

Bulk soil, rhizosphere, and leaf material were homogenized using FastPrep-24™ 5G homogenizer (MP Biomedicals), while roots and grain samples were disrupted with a mortar and liquid nitrogen.
