## Additional File 2_Supplementary Figures for "Plant Growth Promoting fungal endophyte *Colletotrichum tofieldiae* Ct0861 reduces mycotoxigenic *Aspergillus* fungi in maize grains"

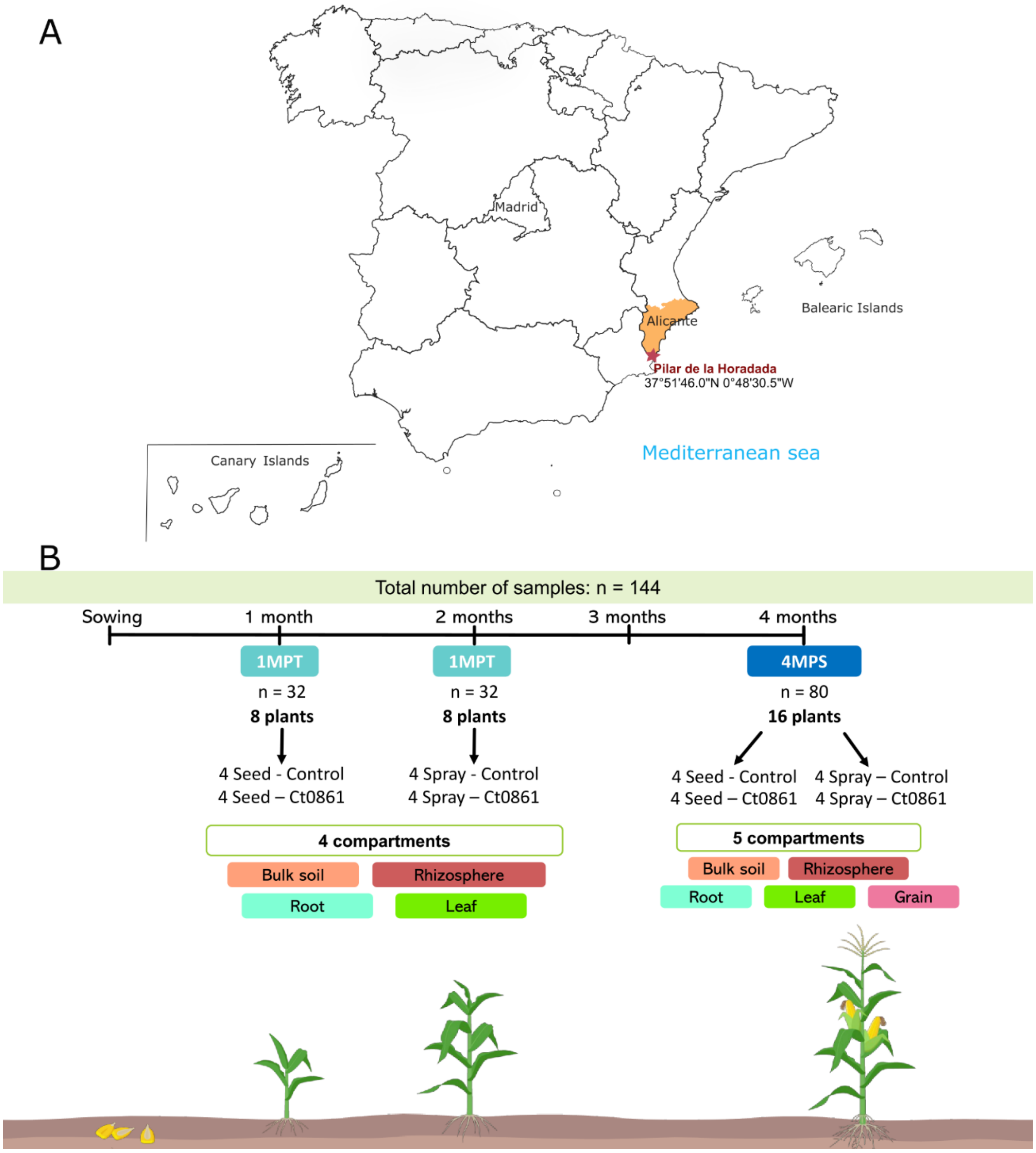


**Supplementary Figure S1:** A) Geographical location of the experimental station. B) Sampling scheme.

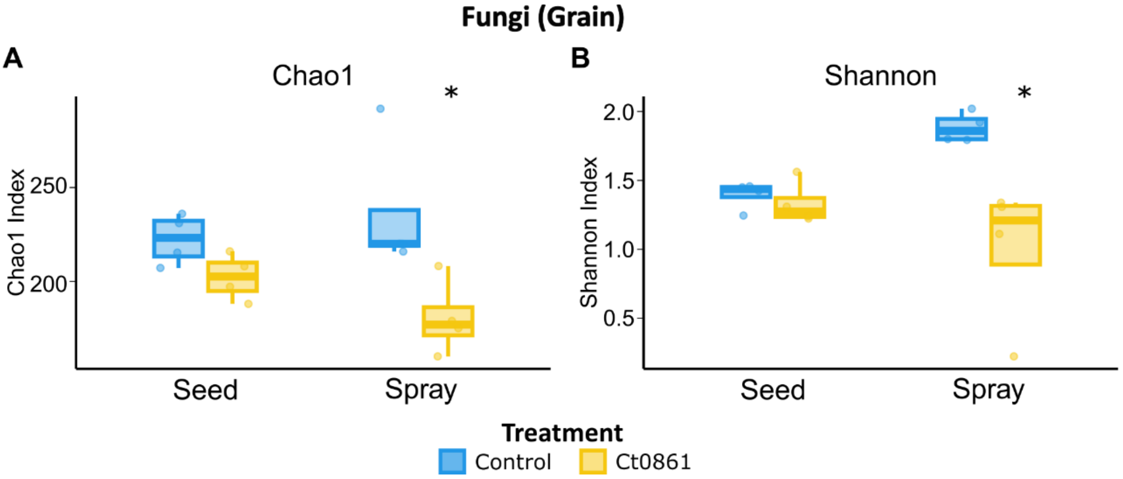


**Supplementary Figure S2: Alpha diversity of grain fungal microbiome treated or not with Ct0861.** Chao1 (A) Shannon indexes (B) of fungal communities in the grain compartment treated or not with Ct0861, either on the seed before sowing or by a spray at one month after sowing. Internal line in box-plot boxes indicates the median or second quartile (Q2), upper line of the boxes indicates the third quartile (Q3) and lower line the first quartile (Q1) of the data. Asterisks denote statistically significant differences in non-parametric Wilcoxon-Mann-Whitney Test considered at $P < 0.05$ (*).

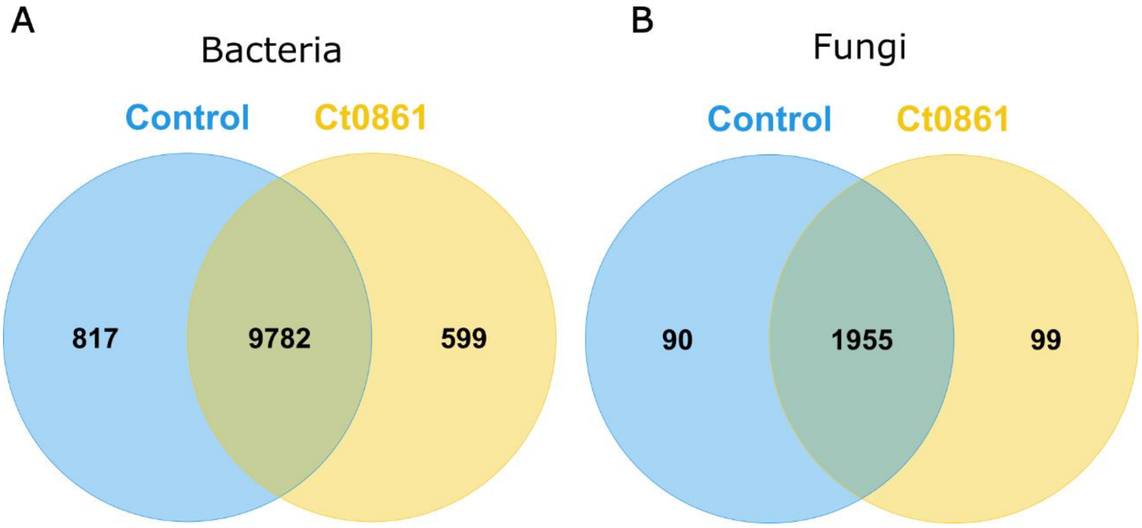


**Supplementary Figure S3.** Venn diagrams of the shared and exclusive (A) bacterial and (B) fungal ASVs between control and Ct0861-treated plants.

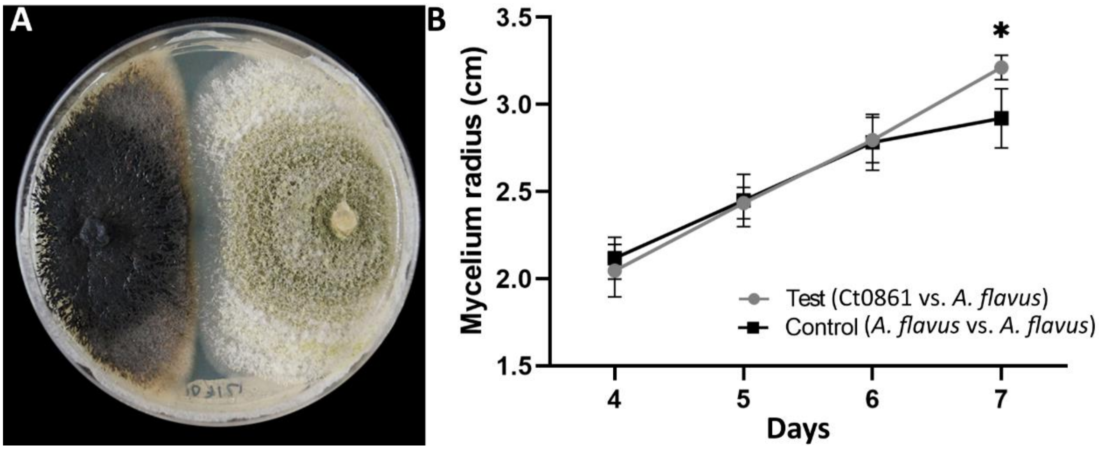


**Supplementary Figure S4: Ct0861-*Aspergillus flavus* dual culture bioassays. A)** Representative image of the cultures after 7 days facing Ct0861 (left) and *A. flavus* NRRL 6540 in PDA plates. B) Mycelial growth of Ct0861-*A. flavus* test and control plates during 7 days. PDA plates facing two plugs of *A. flavus* were used as controls. Values are represented as the average (± standard deviation) of mycelium radius (cm) of each fungus. Asterisks denote statistically significant differences in two-sample Student's T test ($P < 0.05$).
